# The Artists’ Brain: A Data Fusion Approach to Characterize the Neural Bases of Professional Visual Artists

**DOI:** 10.1101/2025.01.01.630982

**Authors:** Erdem Taskiran, Francesca Bacci, David Melcher, Alessandro Grecucci, Nicola De Pisapia

## Abstract

Although everyone has the capacity to draw, only some develop the expertise to produce professional art. Despite extensive creativity research, surprisingly little is known about how years of visual artistic training reshape the neural architecture that distinguishes professional artists from non-artist. To address this gap, we applied joint independent component analysis (jICA) to detect structural (gray matter volume - GM, white matter fractional anisotropy - FA), and functional (resting-state regional homogeneity - ReHo), neuroimaging data from 12 professional visual artists and 12 matched controls. This multimodal approach identified a joint GM-ReHo-FA component (IC2) that significantly distinguished artists from controls (p = .020, d = 1.028). Compared to controls, artists showed coordinated neural adaptations including increased gray matter in parietal, temporal, frontal regions and posterior cingulate cortex; enhanced white matter integrity in anterior thalamic radiations, corticospinal tracts, and association fibers; and increased functional homogeneity in basal ganglia and cerebellar structures. Notably, IC2 expression correlated with higher visual imagery vividness, linking neural adaptations to cognitive abilities fundamental to artistic creation. Taken together, these results highlight the involvement of canonical creativity networks (DMN–ECN) while also extending them to include domain-specific adaptations in cerebellar, sensorimotor, and subcortical systems. Despite these advances, replication with larger samples is necessary.

## Introduction

Artistic creativity is a core aspect of human cognition that enhances the world through music, paintings, design, and other creative outcomes that have influenced cultural evolution (Heilman et al., 2003; Acosta, 2014; Chen et al., 2020). Creativity was traditionally defined by academics like Guilford (1950) as flexibility of mind in generating new and worthy ideas by means of divergent problem-solving. Following Guilford’s early work on divergent thinking—i.e., generating multiple solutions to a problem (Guilford et al., 1967)—our knowledge of creativity has developed considerably. Guilford’s groundbreaking application of divergent thinking set the stage for psychological studies that often approached creativity as a broader cognitive process, while more recent research has highlighted the significance of domain-specific processes, particularly in the arts. This distinction is important: domain-general approaches to creativity focus on cognitive abilities such as fluency and originality that may function across different contexts, whereas domain-specific perspectives emphasize specialized knowledge and techniques unique to particular fields (Baer, 2012; Plucker & Zabelina, 2009).

To better capture the domain-specific nature of artistic practice, the grounded theory of art-making by Mace and Ward (2002) is one of the most useful frameworks. Through extensive interviews and observations with professional artists, they conceptualized artistic creativity as a structured, cyclical process involving planning, problem-finding, idea generation, evaluation, revision, and reflection. These stages are guided by artists’ intentions, aesthetic values, technical skills, and continuous interaction with their developing work. Thus, artistic creativity emerges as a dynamic interaction between intuitive and analytical thinking, deeply rooted in the artist’s identity and practice.

Dissanayake (2001) introduced the concept of Homo Aestheticus, proposing that humans are innately aesthetic beings for whom art-making is a fundamental aspect of our evolutionary heritage. Archaeological evidence, such as beads, ochres, and engravings from sites like Blombos Cave, supports this idea. These artifacts demonstrate that early artistic expression served purposes beyond decoration. It supported social bonding and cultural transmission (Henshilwood et al., 2009). Morriss-Kay (2010) further argued that these symbols functioned similarly to language by reinforcing group identity. Furthermore, recent experimental studies (Rivero et al., 2024) have confirmed that Paleolithic art required advanced visuospatial reasoning and motor control. Together, these perspectives emphasize that artistic creativity is not merely ornamental, but rather an evolutionary adaptation closely linked to cognition and social development.

Building on these psychological and evolutionary perspectives, contemporary neuroscience frames creativity as a complex cognitive process rooted in two key brain networks. The default mode network (DMN)—encompassing medial prefrontal cortex, posterior cingulate cortex, and medial temporal lobes—supports spontaneous ideation and mental simulations essential for divergent thinking (Buckner & DiNicola, 2019; Andrews-Hanna et al., 2010; Mason et al., 2007). The executive control network (ECN)—including lateral prefrontal cortex and posterior parietal cortex—provides cognitive control and evaluative processing necessary to refine ideas into coherent outputs (Seeley et al., 2007; Bressler & Menon, 2010). Neuroimaging studies demonstrate these typically opposing networks co-activate during creative tasks, integrating spontaneous thought with focused evaluation (Beaty et al., 2016, 2019; De Pisapia et al., 2016; Chen et al., 2025). Moreover, individual creative abilities correlate with DMN-ECN connectivity strength (Beaty et al., 2018), while additional regions—anterior cingulate cortex, temporal lobes, and angular gyrus—contribute specialized functions for semantic processing and divergent thinking (Kenett et al., 2023; Jung et al., 2010b).

Beyond the well-established creativity networks, the cerebellum is increasingly recognized as crucial for creative cognition. Although it is traditionally associated with motor control, recent evidence shows that the cerebellum has extensive connections with the cortex that support executive functions and ideational manipulation (Adamaszek et al., 2022; Coolidge, 2021). For example, Saggar et al. (2015) found higher expert-rated creativity correlated with bilateral cerebellar activation, Makuuchi et al. (2003) reported cerebellar involvement in drawing, and Chamberlain et al. (2014) found drawing training increased left cerebellum and SMA volume. Despite this evidence, no studies have examined long-term cerebellar adaptations in professional visual artists

Structural neuroimaging studies have identified positive correlations between creativity and grey matter volumes in multiple regions including frontal, parietal, and dopaminergic areas (Zhu et al., 2013; Sunavsky & Poppenk, 2020; Bendetowicz et al., 2017; Kühn et al., 2014; Takeuchi et al., 2010b; Jung et al., 2013). However, white matter integrity findings are particularly controversial: while some studies link higher fractional anisotropy to better creativity in prefrontal and corticostriatal regions (Takeuchi et al., 2010a; Rahmani et al., 2020), others find inverse relationships, with reduced FA in anterior thalamic radiation and fronto-occipital pathways associated with enhanced creative performance (Jung et al., 2010a; Wertz et al., 2020). Critically, these white matter studies used domain-general cognitive tasks in non-specialist populations. No study has examined how professional artistic training affects white matter integrity, leaving unclear whether these controversial FA-creativity relationships apply to domain-specific artistic expertise.

Further insights come from examining how neurological conditions affect creative behaviors. Research on patients with frontotemporal lobar degeneration (FTLD), Alzheimer’s disease (AD), and Parkinson’s disease (PD) reveals how disruptions in neural pathways impact creativity differently (Acosta, 2014; Miller & Hou, 2004; Schott et al., 2012). FTLD patients often exhibit unexpected artistic behaviors, creating detailed artwork despite cognitive decline, likely due to preserved visuospatial skills in the right parietal cortex (Rankin et al., 2007). However, De Souza et al. (2010) found that frontal variant FTLD patients performed worse on creativity tests, with poor performance linked to frontal pole hypoperfusion. AD patients typically show artistic simplification with diminished detail and muted colors as degeneration affects memory and abstraction regions (Crutch et al., 2001; Piechowski-Jozwiak & Bogousslavsky, 2013). PD patients receiving dopaminergic therapy often show increased creative activity, particularly in divergent thinking and metaphorical creativity. This heightened creativity is linked to elevated dopamine levels enhancing novelty-seeking and ideational fluency (Faust-Socher et al., 2014; Acosta, 2014; Schott, 2012). Professional artists with neurological conditions offer unique insights. Stroke, Alzheimer’s, and aphasia have led to documented artistic transformations—from realistic to abstract styles, altered spatial representation, and shifts in artistic expression that directly reflect underlying neural damage patterns (Crutch et al., 2001; Boller et al., 2005). These clinical cases illustrate how creativity can be transformed by neural change, but leave open the critical question of how the healthy brain adapts over years of professional artistic practice.

Despite extensive creativity research, remarkably few studies have examined professional visual artists with years of specialized training. To our knowledge, only three studies have directly investigated this population. Solso (2001) compared a professional visual artist to a non-artist control during face sketching, observing reduced activation in the right posterior parietal cortex and increased activation in the right middle frontal area in the artist. However, this was a single-subject study. De Pisapia et al. (2016) examined functional connectivity during creative planning tasks, finding that professional visual artists exhibited significantly stronger DMN-ECN connectivity compared to controls. While this aligns with broader creativity findings (Beaty et al., 2019), it captured only task-based functional connectivity. Most recently, Grecucci et al. (2023a) used supervised machine learning to differentiate artists from non-artists based on grey matter features, achieving 79.17% classification accuracy with key regions including Heschl’s gyrus, amygdala and cingulate cortex. Although, classification was successful, the directionality of these grey matter differences remains unclear. Critically, no study has examined how professional artistic expertise is reflected across multiple neuroimaging modalities within the same population. This represents a fundamental gap because creating visual art requires extraordinary coordination between perceptual, motor, and cognitive systems: artists must maintain mental images while manipulating materials, evaluate aesthetic choices while sustaining creative flow, and transform abstract perceptions into tangible forms. These complex demands, refined through thousands of hours of practice, would be expected to leave distinct multimodal signatures that cannot be captured by univariate or unimodal approaches.

### Machine Learning Approach to Understand Artistic Abilities

In recent years, the importance of machine learning techniques in neuroimaging data analysis has grown considerably (Nenning & Langs, 2022). Beyond technical innovation, ML is reshaping how we model brain structure, function, and disease. One primary factor contributing to this popularity is the limitation of mass-univariate methods, which evaluate each voxel individually and fail to consider the statistical interactions present throughout the brain (Baggio et al., 2023). Methods optimized for group-level inference often fail to account for individual variability and distributed network changes, both of which are crucial for comprehending intricate cognitive phenomena such as artistic expertise (Grecucci et al., 2022; Hebart & Baker, 2018; Vieira et al., 2020a). Furthermore, statistical significance of group differences does not ensure separability at the individual level; machine learning redefines success based on predictive accuracy rather than p-values (Schnack, 2020).

Unsupervised learning approaches reveal underlying data structures without predefined target values through clustering and dimensionality reduction (Kherif & Latypova, 2020). Among these methods, Joint Independent Component Analysis (jICA) has emerged as a powerful tool for multimodal neuroimaging fusion. jICA extends ICA principles by concatenating data from different modalities, then decomposing them to identify shared patterns of inter-subject covariation (Calhoun et al., 2006a,b; Adali et al., 2015). This method assumes different imaging techniques may reflect common underlying sources varying consistently across individuals. Unlike hypothesis-driven approaches, jICA’s data-driven nature makes no a priori assumptions, allowing discovery of unexpected patterns (Sui et al., 2012). Given small sample sizes typical of professional artists, jICA is particularly well-suited as it creates simpler models than alternatives (Adali et al., 2015).

We therefore, used jICA to integrate three complementary neuroimaging modalities. Gray matter volume (GM) from structural MRI provides morphometric data associated with artistic abilities (Zhu et al., 2013; Chamberlain et al., 2014; Grecucci et al., 2023a, Li et al., 2019; Shi et al., 2017). Regional homogeneity (ReHo) quantifies local functional synchronization using Kendall’s coefficient of concordance, offering a data-driven measure without requiring predefined ROIs (Zang et al., 2004; Zuo et al., 2013; Lv et al., 2018). From a network perspective, ReHo indicates local network centrality within the functional connectome (Jiang & Zuo, 2016). ReHo has proven valuable in studying creativity (Takeuchi et al., 2017) and serves as a sensitive biomarker of localized brain function (Wang et al., 2011; Xie et al., 2021; Takeuchi et al., 2017). Fractional anisotropy (FA) from diffusion tensor imaging assesses white matter microstructure by measuring water diffusion directionality (Takeuchi et al., 2010a). While higher FA typically associates with better cognitive performance (Bugada et al., 2021; Webb et al., 2020), its role in creativity remains controversial—some studies report positive associations (Takeuchi et al., 2010a; Rahmani et al., 2020), others find inverse relationships (Jung et al., 2010a; Wertz et al., 2020).

The fundamental advantage of jICA lies in accessing joint information across modalities that cannot be obtained from separate analyses (Sui et al., 2012; Calhoun & Sui, 2016). By identifying components that covary across three neuroimaging modalities, jICA can reveal coordinated adaptations reflecting years of artistic training (Lahat et al., 2015). Multimodal fusion approaches have successfully identified distributed neural signatures in various psychological phenomena, including emotional intelligence and anxiety (Grecucci et al., 2024), borderline personality disorder (Grecucci et al., 2023b), phobic disorders (Scarano et al., 2025) and many more (See Grecucci et al., 2025). By pooling evidence across modalities, multimodal analyses increase statistical power to detect weaker or more dispersed patterns— crucial given the typically small samples of professional artists available for neuroimaging research.

### Current Study and Hypotheses

Building upon this theoretical and methodological foundation, our primary hypothesis is that professional visual artists will exhibit distinct multimodal neural signatures characterized by joint independent components integrating structural, and functional differences across distributed brain networks. We predict these signatures will encompass both domain-general creativity networks (DMN and ECN) and domain-specific adaptations unique to visual artistic expertise. Supporting this prediction, Gupta et al. (2019) demonstrated that ICA-based decompositions of structural data often yield components corresponding to canonical functional networks including the DMN, ECN, and salience networks, reinforcing the biological validity of targeting these systems through multimodal analysis. Furthermore, Vessel and colleagues’ fMRI studies reveal that DMN regions are selectively engaged during emotionally powerful aesthetic experiences (Vessel et al., 2012, 2013, 2019), raising the question of whether years of artistic training produce corresponding structural adaptations in these functionally-relevant regions. We expect increased structural density in regions overlapping with these canonical networks, consistent with Kühn et al.’s (2014) findings of structural DMN correlates in creativity. However, we anticipate these patterns will extend beyond traditional creativity networks, as domain-general and domain-specific creativity share some neural substrates while maintaining distinct features (Chen et al., 2020).

Based on growing evidence for cerebellar involvement in creative cognition (Adamaszek et al., 2022; Coolidge, 2021; Cotterill, 2001; Mumford & Caughron, 2007; Gao et al., 2020), we hypothesized enhanced cerebellar representation across modalities in professional artists. This may reflect the cerebellum’s dual role in motor control for precise artistic execution and cognitive processes supporting creative ideation through internal model generation (Ito, 2008). Within the predictive coding framework (Frascaroli et al., 2024), the cerebellum’s capacity for generating and updating internal models could serve as the neural foundation for predictive mechanisms supporting artistic creativity.

Given the controversial FA findings in creativity literature, we predict specific white matter alterations in: cerebellar-cortical connections, frontal executive control tracts, and visuomotor pathways. Additionally, considering the physical demands of artistic practice, we expect enhanced representation in sensorimotor areas supporting precise hand movements and sensory feedback (Chamberlain et al., 2014), along with parietal contributions supporting visuospatial processing (Kozbelt & Seeley, 2007).

## Materials and Methods

### Participants

The study involved a total of 24 participants (14 males and 10 females), consisting of 12 professional visual artists and 12 non-artists. All participants were right-handed and reported no history of psychiatric or neurological disorders, nor any use of drugs affecting the central nervous system. The professional artists were recruited with the collaboration of the Museum of Modern and Contemporary Art (MART) in Rovereto, Italy. Their professional status was confirmed through institutional recognition within the Italian art system. These artists also have established careers, as evidenced by their participation in significant exhibitions and their active engagement in the professional art community. Nine of the artists received formal training in art schools or academies, where they studied the fundamental visual elements-such as color, form, line, space, texture, and value-and developed expertise in techniques such as sketching, drawing, and painting. This training included the mastery of proportion, shading, and three-dimensional modeling. The remaining three artists were self-taught, having independently cultivated their skills despite lacking formal academic instruction. The control group comprised individuals whose professions and skills were unrelated to drawing or visual arts. They were recruited through local advertisements and were screened prior to participation to ensure that none were professional artists or engaged in visual artistic activities. Additionally, all individuals were asked to create a drawing immediately after the scanning session. This procedure not only supported the aims of the original study but also served to assess the artistic ability of each participant. The works created by the artists were later developed into completed pieces and featured in a curated museum exhibition titled “In Risonanza: Istantanee di Creatività nel Cervello” held at MART (Rovereto, Italy) (Bacci et al., 2013). This exhibition publicly displayed the creative outputs of the artists and offered further confirmation of their professional status. For more information about this exhibition, see: https://www.mart.tn.it/mostre/in-risonanza-istantanee-di-creativita-nel-cervello-138701, and refer to the original publication (De Pisapia et al., 2016). The two groups were closely matched for age and gender. The mean age of the artists was 30.9 years, while that of the non-artists was 29.78 years (p = 0.59). Gender distribution was comparable between the groups, with the artist group consisting of 4 females and 8 males, and the control group comprising 6 females and 6 males (p = 0.41) (see Table 1. Below and table 4 in supplementary material for participant specific information). All procedures were conducted in accordance with relevant guidelines and regulations and received approval from the University of Trento Ethics Committee (Protocol Number: 2008-006).

**Table 1.** Sample Information.

| Variable | Artists(n=12) | Control(n=12) | Statistics | p-Value |
| --- | --- | --- | --- | --- |
| Age (years) | 30.90(+/-4.76) | 29.77 (+/- 5.32) | $t=0.551$ | 0.587 |
| Gender(M/F) | 8/4 | 6/6 | $\chi^2= 0.686$ | 0.408 |
| VVIQ Score | 29.18 (+/- 7.07) | 36.50 (+/- 6.86) | $t=-2.520$ | 0.020 |

### Behavioral Assessment: Vividness of Visual Imagery Questionnaire (VVIQ)

All participants completed the Vividness of Visual Imagery Questionnaire (VVIQ), a widely used self-report measure designed to assess the clarity and vividness of internally generated visual images (Marks, 1973). The questionnaire consists of 16 items, each prompting individuals to imagine a specific scenario (e.g., a relative’s face, a sunset) and rate the vividness of their mental imagery on a five-point Likert scale. A score of 1 indicates an image “perfectly clear and as vivid as normal vision,” whereas a score of 5 reflects “no image at all—you only know that you are thinking of the object.” *Lower scores thus correspond to greater imagery vividness*. Original version of the VVIQ was selected, rather than the revised VVIQ-2 (Marks, 1995), due to its extensive validation, structural comparability to the updated version, and practical advantages; such as reduced administration time and cognitive demand, especially given the breadth of current imaging and behavioral protocol.

### Data Acquisition

All participants underwent magnetic resonance imaging at the Center for Mind Brain Sciences (CIMeC), following screening for MRI compatibility by a medical professional. The imaging protocol was performed on a 4 Tesla Siemens MedSpec scanner system and encompassed structural T1-weighted imaging, resting-state functional MRI, and diffusion-weighted imaging in a single session. Participants’ heads were stabilized using foam padding throughout the acquisition to minimize motion artifacts.

High-resolution anatomical images were acquired using a birdcage transmit coil and an 8-channel radio-frequency phased-array head receiver coil. T1-weighted images were obtained using a three-dimensional magnetization-prepared rapid gradient-echo (MP-RAGE) sequence with the following parameters: voxel resolution = 1 x 1 x 1 mm³, field of view = 256 x 224 mm², 176 contiguous sagittal slices, GRAPPA parallel imaging acceleration factor = 2, repetition time = 2700 ms, echo time = 4.18 ms, inversion time = 1020 ms, and flip angle = 7°. The structural imaging required approximately 6 minutes of acquisition time.

Resting-state functional MRI data were collected using a gradient-echo echo-planar imaging sequence with voxel resolution = 3 x 3 x 3 mm³, TR = 2200 ms, echo time = 33 ms, flip angle = 75°, 37 axial slices acquired in interleaved ascending order with 3 mm thickness covering the whole brain. The functional acquisitions utilized anterior-posterior phase encoding direction with an echo spacing of 0.3392 ms and total readout time of 21.37 ms, without parallel imaging acceleration.

Diffusion-weighted imaging data were acquired using a single-shot spin-echo echo-planar imaging (SE-EPI) sequence. The acquisition protocol employed the following parameters: repetition time (TR) = 7000 ms, echo time (TE) = 85 ms, flip angle = 90°, acquisition matrix = 96 x 96, field of view = 240 x 240 mm², voxel size = 2.5 x 2.5 x 2.5 mm³, 54 contiguous axial slices with no interslice gap, bandwidth = 2365 Hz/pixel, and GRAPPA parallel imaging acceleration factor = 2. The diffusion-weighting scheme comprised 35 volumes in total, consisting of 5 non-diffusion-weighted volumes (b = 0 s/mm²) acquired at the beginning of the sequence, followed by 30 diffusion-weighted volumes (b = 1000 s/mm²) with gradient directions uniformly distributed on the sphere. All diffusion acquisitions utilized anterior-posterior phase encoding direction (j-) with fat saturation enabled. Signal reception was performed using an 8-channel phased-array head coil.

### Pre-Processing Steps

#### Structural Grey Matter Pre-processing

All T1-weighted structural images underwent a quality assessment to exclude any artifacts prior to analysis. Preprocessing was performed using SPM12 (Statistical Parametric Mapping; https://www.fil.ion.ucl.ac.uk/) (Penny et al., 2011) in conjunction with the Computational Anatomy Toolbox (CAT12; http://www.neuro.uni-jena.de/cat/) (Gaser et al., 2024) within the MATLAB environment. Initially, each image was re-oriented to position the anterior commissure at the origin. Following re-orientation, the images were segmented into gray matter (GM), white matter (WM), and cerebrospinal fluid (CSF) using CAT12. The analysis exclusively utilized gray matter (GM) images to the application of Joint Independent Component Analysis. Subsequent registration was conducted using Diffeomorphic Anatomical Registration through Exponential Lie algebra tools (Ashburner, 2007), which facilitates a whole-brain approach to align anatomical structures across participants. After registration, the DARTEL-normalized images were transformed to the Montreal Neurological Institute (MNI) space. To prepare the data for further analysis, a spatial Gaussian smoothing with a kernel size of 10 mm, was applied.

#### RS-fMRI Pre-processing

Data preprocessing was performed using the Data Processing Assistant for Resting-State fMRI (DPARSF) V5.3 Advanced Edition (State Key Laboratory of Cognitive Neuroscience and Learning, Beijing Normal University, China) (Yan & Zang, 2010), which is based on the Data Processing and Analysis of Brain Imaging (DPABI) Toolbox version 8.2 (Yan et al., 2016) with statistical parametrical mapping 12 (SPM12; Wellcome Trust Center for Neuroimaging, University College London, UK) in Matlab 2023a (MathWorks, Inc., Natick, MA, USA).

The first 5 volumes were discarded and the subsequent 215 volumes underwent slice timing correction using the middle slice as the reference slice, followed by realignment for head motion correction. Following strict quality control protocols, participants were excluded if their head motion exceeded 2.0 mm of maximal translation in any direction (x, y, or z) or 2.0° of maximal rotation during scanning (Wang et al., 2015; Di & Biswal, 2023; Yuan et al., 2018). No subject in each group was excluded from further analysis based on these criteria.

The spatial normalization process involved several steps. Initially, each image was re-oriented to position the anterior commissure at the origin manually, and secondly, brain extraction was performed using FSL’s Brain Extraction Tool (BET) (Smith, 2002) within the DPARSF (Yan & Zang, 2010) framework. Following brain extraction, individual T1 structural images were co-registered to their corresponding functional images. Subsequently, these co-registered T1 images were segmented via DARTEL (Ashburner, J. 2007; Ashburner & Friston, 2005).

Finally, all functional images were transformed into Montreal Neurological Institute (MNI) standard space and resampled to 3 x 3 x 3 mm³ voxels (Yang et al., 2021). To minimize the influence of nuisance signals, we implemented the Friston 24-parameter model (Friston et al., 1996), which includes six head motion parameters, six head motion parameters from one time point before, and their twelve corresponding squared items. Additionally, signals from white matter, cerebrospinal fluid, and global signals were regressed out. Global signal removal has been shown to reduce physiological noise and movement-related effects (Yan et al., 2013).

Linear trend removal was performed, followed by temporal band-pass filtering (0.01-0.08 Hz) (Satterthwaite et al., 2013). After preprocessing, the quality control was performed according to the standardized procedures outlined by Lu and Yan (2023) in the DPABI framework. No participants were excluded based on this quality control criteria.

For Regional Homogeneity (ReHo) calculation (Zang et al., 2004), we implemented a specific preprocessing stream in DPARSF based on the fundamental principle that fMRI activity is more likely to occur in clusters of several spatially contiguous voxels than in a single voxel (Tononi et al., 1998; Katanoda et al., 2002; Ganos et al., 2014). Importantly, spatial smoothing was performed after, not before the ReHo calculation to prevent artificial inflation of local synchronization (Zang et al., 2004; Yan & Zang, 2010). For each voxel, Kendall’s coefficient of concordance (KCC) (Kendall & Gibbons, 1990) was calculated between its time series and those of its 26 nearest neighbors. The individual ReHo maps were then standardized by dividing by the global mean KCC within the whole-brain mask followed by spatial smoothing with a 6 mm FWHM Gaussian kernel. Following computation, ReHO maps underwent standardization for group-level analysis. Z-scores were calculated by subtracting the mean value and dividing by the standard deviation. This standardization enables meaningful interpretation where positive Z-scores indicates a higher regional synchronization (ReHO) in that individual’s brain, while negative Z-scores indicate lower regional synchronization.

Individual z-transformed ReHO maps were used for further fusion analysis.

### Preprocessing Pipeline for DTI

All diffusion-weighted imaging data underwent comprehensive preprocessing using the FMRIB Software Library (FSL version 6.0; Jenkinson et al., 2012). The preprocessing pipeline was an automated bash script created by the Authors (See Supplementary Material for the Github Link) to ensure consistency across all subjects. Initially, raw DICOM images were converted to NIfTI format using dcm2niix (Li et al., 2016). The first non-diffusion-weighted volume (b = 0 s/mm²) was extracted from the complete dataset. Brain extraction was subsequently performed on this b0 volume using the Brain Extraction Tool (BET; Smith, 2002).

Given the absence of reverse phase-encoded acquisitions in the current dataset, susceptibility-induced distortion correction using the TOPUP algorithm (Andersson et al., 2003) was not feasible. However, comprehensive correction for eddy current-induced distortions and subject motion was performed using FSL’s eddy tool (Andersson & Sotiropoulos, 2016). Following distortion and motion correction, diffusion tensors were fitted to the preprocessed data. The tensor fitting procedure generated multiple parametric maps and only FA maps used for the purpose of this study. Spatial normalization of the FA maps to standard space was achieved through a two-stage registration procedure. First, affine registration was performed using FLIRT (FMRIB’s Linear Image Registration Tool; Jenkinson & Smith, 2001), aligning each subject’s FA map to the FMRIB58_FA template in MNI152 space at 1 x 1 x 1 mm³ resolution. For the subsequent non-linear registration, we used FNIRT (FMRIB’s Non-linear Image Registration Tool; Andersson et al., 2007). Finally, the normalized FA maps underwent spatial smoothing using a Gaussian kernel with full width at half maximum (FWHM) of 8 mm to increase signal-to-noise ratio. Some parts of the DTI preprocessing steps were adapted based on the procedures described in Kim et al. (2015) in their mCCA+jICA (GM+FA) fusion study. Comprehensive quality assessment was performed at both individual and group levels. Individual subject data quality was evaluated using eddy_quad (Bastiani et al., 2019).

Subsequently, group-level quality assessment was conducted using eddy_squad (Bastiani et al., 2019). Based on these quality metrics, all 24 subjects (FA maps) met the inclusion criteria and were retained for subsequent statistical analyses.

### Multimodal Fusion Analysis: Joint Independent Component Analysis (jICA)

Joint Independent Component Analysis (jICA) represents a powerful data-driven approach for identifying linked patterns across multiple neuroimaging modalities that share common sources of intersubject covariation (Calhoun et al., 2006a, b) (See Figure 2.). jICA is based on a strong assumption that different modalities can be fused to yield the same mixing coefficient matrix (Calhoun et al., 2006a; Yang et al., 2019). The mathematical framework of jICA assumes a generative model where the observed multimodal data matrix ***X*** can be decomposed as the product of a mixing matrix **A** and a source matrix ***S***, mathematically expressed as ***X*** *= **AS***. In this formulation, the joint observation matrix ***X*** *∈ ℝ^(N×V)* is constructed through horizontal concatenation of the normalized modality-specific data matrices: ***X*** *= [**X**^(GM)^ | **X**^(ReHo)^] | **^X(^**^FA)^]* where N = 24 represents the number of subjects and V represents the total number of voxels across all three modalities after brain masking (Franco et al., 2008). The mixing matrix ***A*** contains the subject-specific loading coefficients that indicate the degree to which each independent component represents a subject’s multimodal data as a whole (Lerman-Sinkoff et al., 2017), while the source matrix ***S*** comprises K spatially independent joint sources that can be partitioned back into modality-specific maps. Prior to ICA decomposition, dimensionality reduction was performed via principal component analysis (PCA) to enhance computational efficiency while preserving the essential variance structure of the data (Calhoun et al., 2006a, b). This data reduction step, implemented through singular value decomposition (SVD) (Lahat et al., 2015), retained components that explained %93.58 of the total variance across modalities. The optimal number of independent components was estimated using the modified minimum description length (MDL) criterion (Li et al., 2007). The MDL analysis yielded different optimal component numbers for each modality: 5 for GM,7 for ReHo and 6 for FA. Following the recommendation of Sui et al. (2011), who demonstrated through simulation that slight overestimation of component number does not adversely affect results, we therefore selected K = 7 components for the joint analysis. The joint ICA decomposition was performed using the FastICA algorithm (Hyvärinen, 1999; Hyvärinen & Oja, 2000), which employs a fixed-point iteration scheme to maximize the non-Gaussianity of the estimated sources. To address the stochastic nature of ICA initialization, we implemented the ICASSO Algorithm (Himberg et al., 2004) with 10 runs, clustering the resulting components across iterations and computing stability indices (Iq) for each component. Components demonstrating stability indices exceeding 0.8 were retained. Joint ICA Analysis was performed using the Fusion ICA Toolbox (FIT v2.0.5.4; https://github.com/trendscenter/fit )in conjunction with the Group ICA Toolbox (GIFT v4.0.6.5; https://github.com/trendscenter/gift) implemented in MATLAB 2023a.

**Figure 1.**
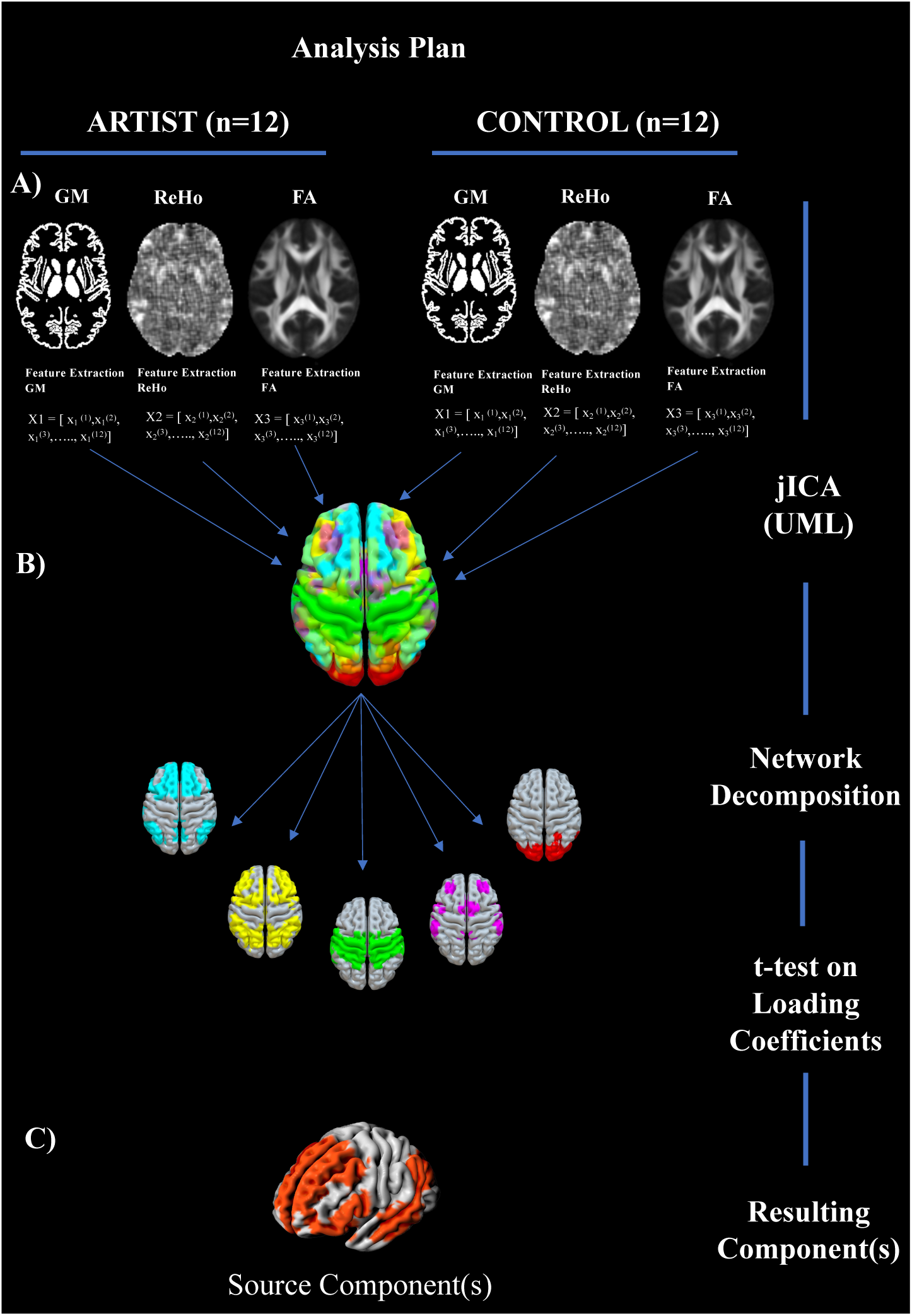
Overview of the joint ICA (jICA) multimodal fusion pipeline**. (A)** For each participant (12 artists, 12 controls), modality-specific brain features were extracted: gray matter (GM), regional homogeneity (ReHo) and fractional anisotropy (FA). Each participant’s data for each modality was reshaped into a one-dimensional vector of non-zero voxels**. (B)** The resulting matrices from GM, ReHo, and FA were normalized and concatenated across participants to form the input for joint ICA. The jICA decomposition produced a set of spatially independent components (joint sources), each shared across modalities (K=7). These components were visualized as brain maps per modality**. (C)** A subset of components that significantly differentiated groups (artists vs. controls) based on their mixing coefficients was further visualized as 3D source maps.

**Figure 2.**
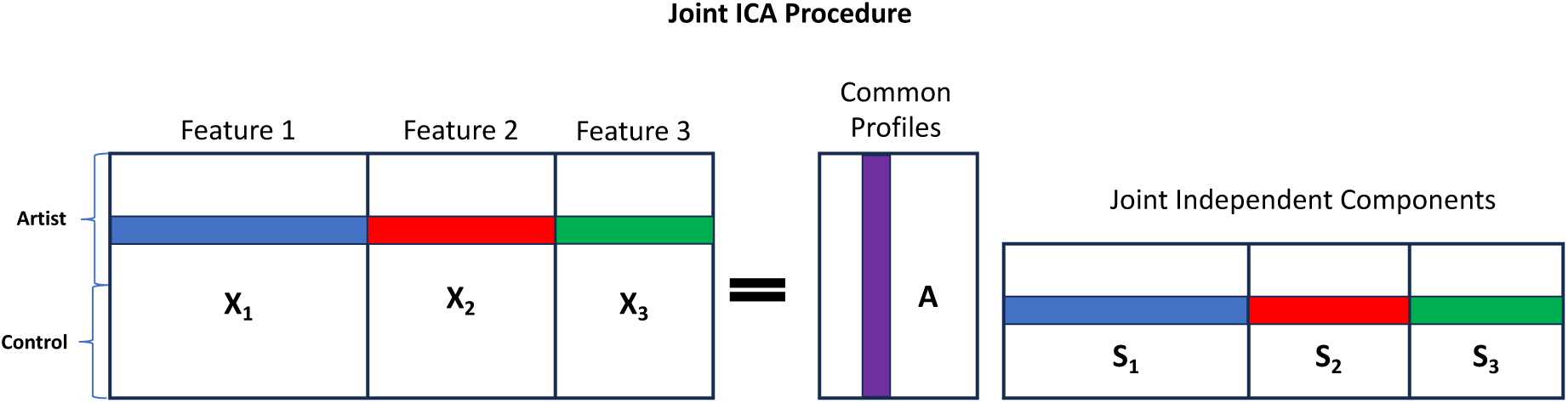
Overview of the joint ICA (jICA) multimodal fusion procedure. The input matrix (left) consists of voxel-wise feature matrices for gray matter (GM), regional homogeneity (ReHo), and fractional anisotropy (FA), horizontally concatenated across participants (12 artists and 12 controls). This results in a joint data matrix, X = [X_1_, X_2_, X_3_], where each row represents a subject and each column represents a feature from one of the three modalities. Joint independent component analysis (jICA) is then performed on the matrix, yielding a shared mixing matrix A that captures individual expression levels for each component (middle) and a source matrix S = [S_1_, S_2_, S_3_] containing modality-specific spatial maps (right) (The figure inspired by (Wang et al., 2015)).

### Statistical Analysis of Mixing Coefficients

The output of the joint ICA decomposition includes the mixing matrix **A** (24 x 7), which contains the subject-specific mixing coefficients or ICA loadings (See Figure 1. For overview). These loading parameters serve as a compact representation of how strongly each joint source, comprising linked patterns of gray matter volume, functional homogeneity, and white matter integrity, is expressed in each individual. Conceptually, higher absolute values indicate greater expression of that particular multimodal pattern, while the sign (+/-) indicates the direction of this expression relative to the group mean.

To identify joint components that differentiate professional visual artists from non-artists, independent samples Student’s t-tests were conducted on the mixing coefficients of each component. For components that did not meet the assumption of normality, as assessed by the Shapiro-Wilk test (α = 0.05), the non-parametric Mann-Whitney U test was applied.

After conducting group comparisons, we investigated whether the multimodal components that had significantly differentiated between professional visual artists and non-artists were also associated with visual imagery abilities. Specifically, we tested for correlations between the mixing coefficients of these components and participants’ scores on the Vividness of Visual Imagery Questionnaire (VVIQ). This allowed us to determine whether the neural patterns that distinguish artists from non-artists also reflect differences in the vividness of mental imagery. All statistical analyses were performed using JASP (version 0.19.2; 2025).

## RESULTS

### Group Differences in Joint Component Expression

After the normalization (see Supplementary material for detailed Normalization step prior to jICA) Joint ICA decomposition was applied across three modalities, yielding seven joint sources, as determined by the minimum description length (MDL) approach (Li et al., 2007). An independent samples two-tailed Student’s *t*-test was conducted on the mixing coefficients (***A***) to assess between-group differences. Among the seven joint sources, only IC2 showed a statistically significant difference between artists and controls (Figure 3): *t*(22) = 2.517, *p* = .020, Cohen’s *d* = 1.028. This indicates a large effect size (Cohen, 2013), with artists (M = 0.177, SD = 0.090) exhibiting greater expression of IC2 compared to controls (M = 0.092, SD = 0.075). The Shapiro–Wilk test confirmed that residuals for IC2 were normally distributed (*W* = 0.957, *p* = .387), and the Brown–Forsythe test indicated homogeneity of variances (*F* = 0.678, *p* = .419), satisfying t-test assumptions. No significant group differences were found for the remaining components (all *ps*> .15). There was no significant correlation between age and IC2 scores (*r* = –0.171, *p* = .424), and no significant gender effect on IC2 loadings (*t*(22) = 0.566, *p* = .577). Spatial maps were thresholded at Z > 4 for gray matter and Z > 3.5 for both fractional anisotropy and regional homogeneity (see supplementary materials for detailed visualization step). Anatomical labeling was performed using Talairach coordinates for gray matter and ReHo maps (see Table 2), while JHU ICBM-DTI-81 20-Tract Atlas was used for FA tract identification (see Table 3 and see Supplementary material for the detailed FA tract identification).

**Figure 3.**
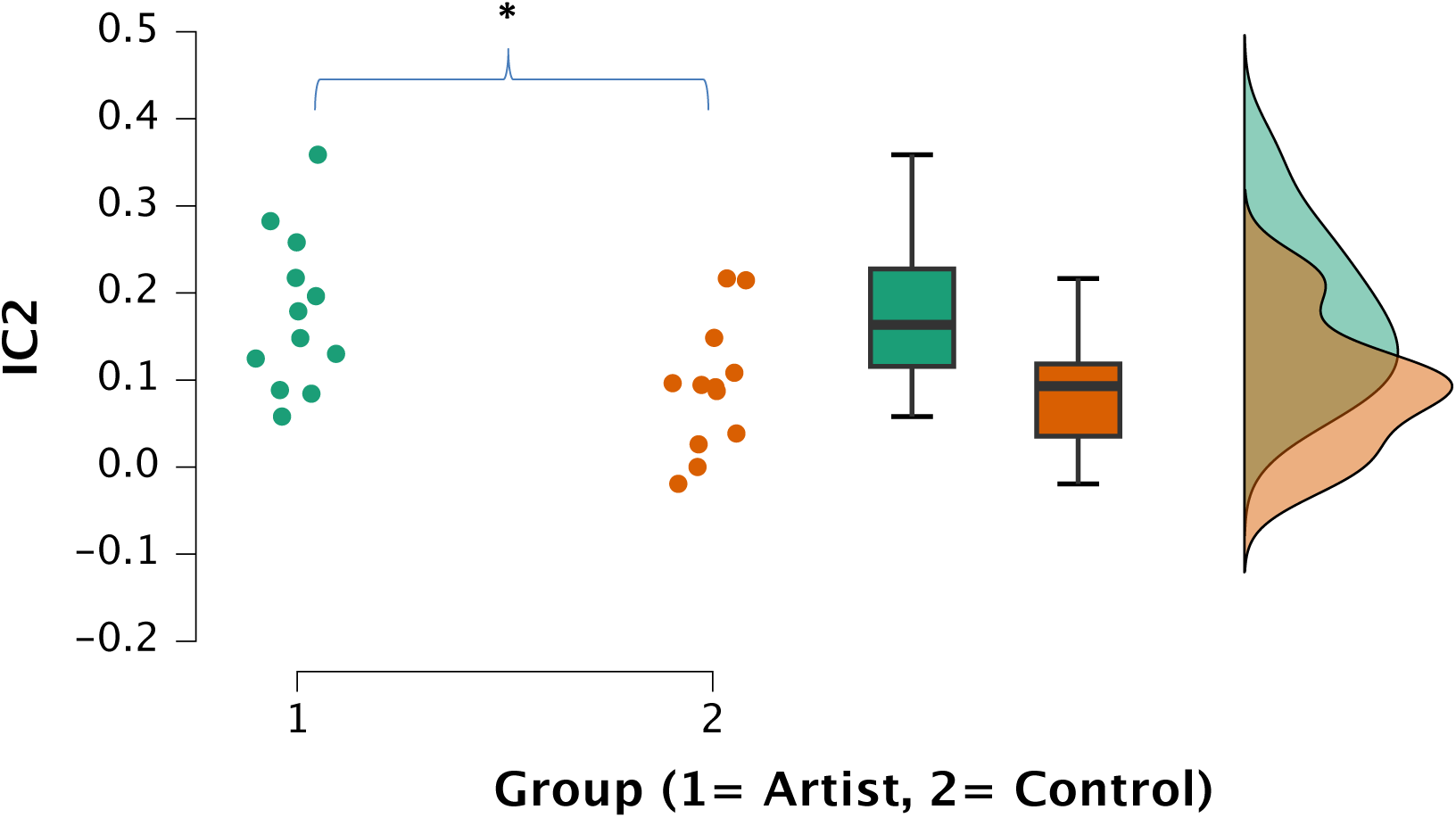
Group differences in IC2 loading coefficients. Artists (Group 1) exhibited significantly greater expression of IC2 (*M* = 0.177, *SD* = 0.090) than controls (Group 2; *M* = 0.092, *SD* = 0.075), *t*(22) = 2.517, *p* = .020, Cohen’s *d* = 1.028

**Figure 4.**
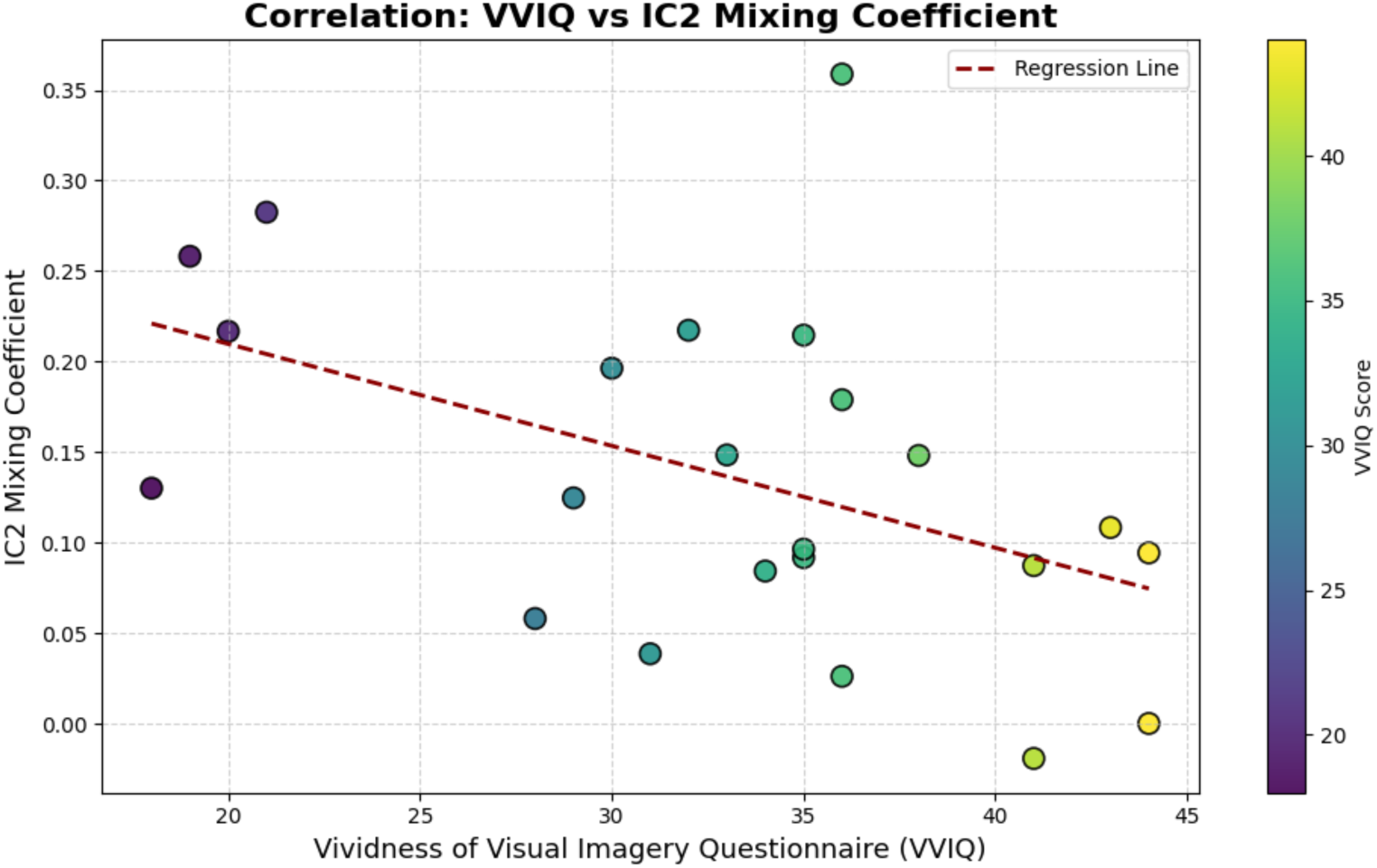
The scatter plot showing the negative correlation between IC2 mixing coefficients and VVIQ total scores. Each dot represents a participant and is coloured according to their VVIQ score (higher values indicate less vivid imagery scores).

**Table 2.**
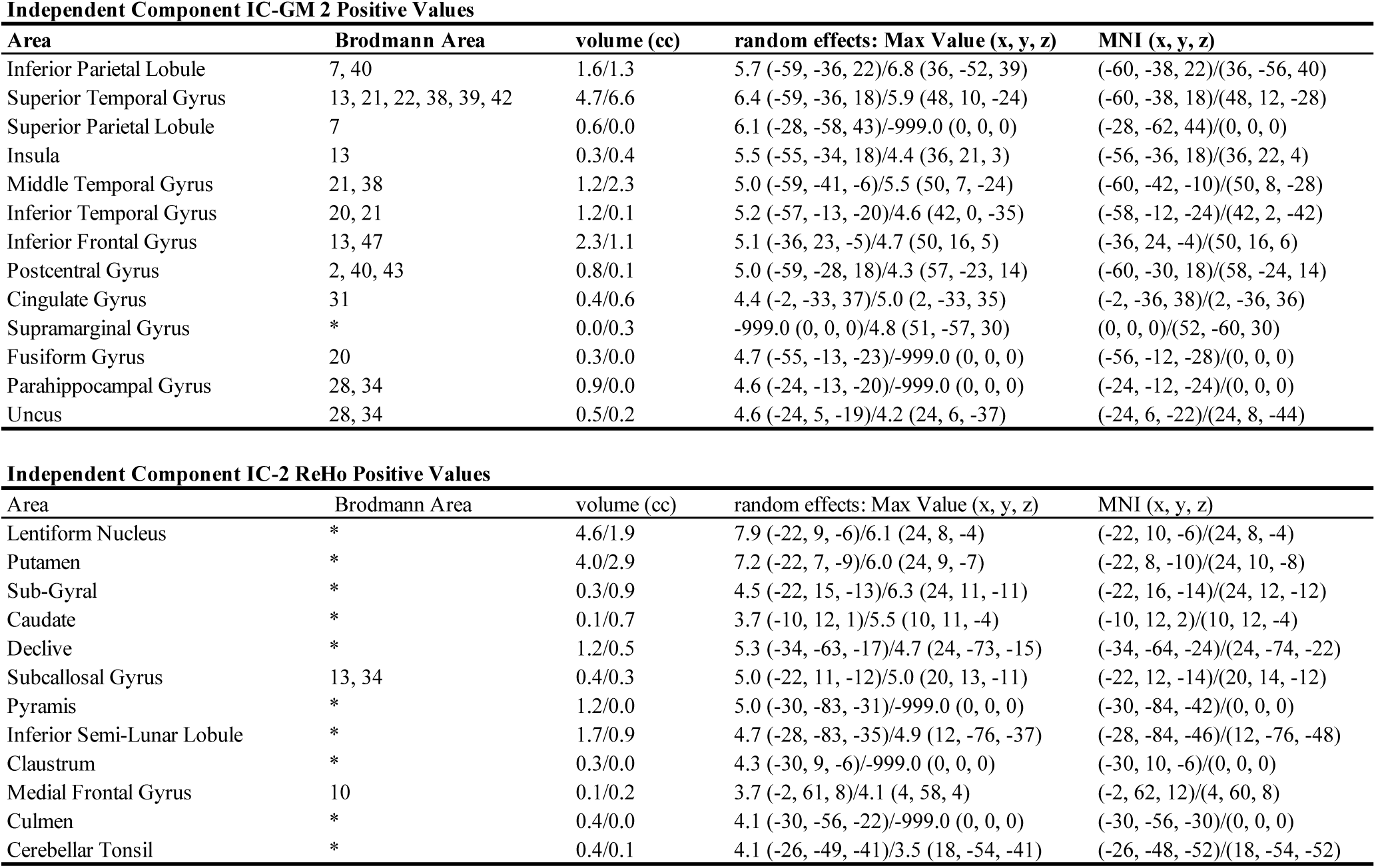
IC2 GM and ReHo Positive Regions.

| Independent Component IC-GM 2 Positive Values |  |  |  |  |
| --- | --- | --- | --- | --- |
| Area | Brodman Area | volume (cc) | random effects: Max Value (x, y, z) | MNI (x, y, z) |
| Inferior Parietal Lobule | 7, 40 | 1.6/1.3 | 5.7 (-59, -36, 22)/6.8 (36, -52, 39) | (-60, -38, 22)/(36, -56, 40) |
| Superior Temporal Gyrus | 13, 21, 22, 38, 39, 42 | 4.7/6.6 | 6.4 (-59, -36, 18)/5.9 (48, 10, -24) | (-60, -38, 18)/(48, 12, -28) |
| Superior Parietal Lobule | 7 | 0.6/0.0 | 6.1 (-28, -58, 43)/-999.0 (0, 0, 0) | (-28, -62, 44)/(0, 0, 0) |
| Insula | 13 | 0.3/0.4 | 5.5 (-55, -34, 18)/4.4 (36, 21, 3) | (-56, -36, 18)/(36, 22, 4) |
| Middle Temporal Gyrus | 21, 38 | 1.2/2.3 | 5.0 (-59, -41, -6)/5.5 (50, 7, -24) | (-60, -42, -10)/(50, 8, -28) |
| Inferior Temporal Gyrus | 20, 21 | 1.2/0.1 | 5.2 (-57, -13, -20)/4.6 (42, 0, -35) | (-58, -12, -24)/(42, 2, -42) |
| Inferior Frontal Gyrus | 13, 47 | 2.3/1.1 | 5.1 (-36, 23, -5)/4.7 (50, 16, 5) | (-36, 24, -4)/(50, 16, 6) |
| Postcentral Gyrus | 2, 40, 43 | 0.8/0.1 | 5.0 (-59, -28, 18)/4.3 (57, -23, 14) | (-60, -30, 18)/(58, -24, 14) |
| Cingulate Gyrus | 31 | 0.4/0.6 | 4.4 (-2, -33, 37)/5.0 (2, -33, 35) | (-2, -36, 38)/(2, -36, 36) |
| Supramarginal Gyrus | * | 0.0/0.3 | -999.0 (0, 0, 0)/4.8 (51, -57, 30) | (0, 0, 0)/(52, -60, 30) |
| Fusiform Gyrus | 20 | 0.3/0.0 | 4.7 (-55, -13, -23)/-999.0 (0, 0, 0) | (-56, -12, -28)/(0, 0, 0) |
| Parahippocampal Gyrus | 28, 34 | 0.9/0.0 | 4.6 (-24, -13, -20)/-999.0 (0, 0, 0) | (-24, -12, -24)/(0, 0, 0) |
| Uncus | 28, 34 | 0.5/0.2 | 4.6 (-24, 5, -19)/4.2 (24, 6, -37) | (-24, 6, -22)/(24, 8, -44) |

| Independent Component IC-2 ReHo Positive Values |  |  |  |  |
| --- | --- | --- | --- | --- |
| Area | Brodman Area | volume (cc) | random effects: Max Value (x, y, z) | MNI (x, y, z) |
| Lentiform Nucleus | * | 4.6/1.9 | 7.9 (-22, 9, -6)/6.1 (24, 8, -4) | (-22, 10, -6)/(24, 8, -4) |
| Putamen | * | 4.0/2.9 | 7.2 (-22, 7, -9)/6.0 (24, 9, -7) | (-22, 8, -10)/(24, 10, -8) |
| Sub-Gyral | * | 0.3/0.9 | 4.5 (-22, 15, -13)/6.3 (24, 11, -11) | (-22, 16, -14)/(24, 12, -12) |
| Caudate | * | 0.1/0.7 | 3.7 (-10, 12, 1)/5.5 (10, 11, -4) | (-10, 12, 2)/(10, 12, -4) |
| Declive | * | 1.2/0.5 | 5.3 (-34, -63, -17)/4.7 (24, -73, -15) | (-34, -64, -24)/(24, -74, -22) |
| Subcallosal Gyrus | 13, 34 | 0.4/0.3 | 5.0 (-22, 11, -12)/5.0 (20, 13, -11) | (-22, 12, -14)/(20, 14, -12) |
| Pyramis | * | 1.2/0.0 | 5.0 (-30, -83, -31)/-999.0 (0, 0, 0) | (-30, -84, -42)/(0, 0, 0) |
| Inferior Semi-Lunar Lobule | * | 1.7/0.9 | 4.7 (-28, -83, -35)/4.9 (12, -76, -37) | (-28, -84, -46)/(12, -76, -48) |
| Clastrum | * | 0.3/0.0 | 4.3 (-30, 9, -6)/-999.0 (0, 0, 0) | (-30, 10, -6)/(0, 0, 0) |
| Medial Frontal Gyrus | 10 | 0.1/0.2 | 3.7 (-2, 61, 8)/4.1 (4, 58, 4) | (-2, 62, 12)/(4, 60, 8) |
| Culmen | * | 0.4/0.0 | 4.1 (-30, -56, -22)/-999.0 (0, 0, 0) | (-30, -56, -30)/(0, 0, 0) |
| Cerebellar Tonsil | * | 0.4/0.1 | 4.1 (-26, -49, -41)/3.5 (18, -54, -41) | (-26, -48, -52)/(18, -54, -52) |

**Table 3.** IC2-FA Positive Tracts.

| Independent Component IC-FA 2 Positive Values |  |  |  |  |  |  |
| --- | --- | --- | --- | --- | --- | --- |
| TractName | Voxels | Volume (cc) | Max Z | X | Y | Z |
| Anterior thalamic radiation L | 128 | 1.0 | 7.381425 | 50.835938 | 40.695312 | 15.328125 |
| Anterior thalamic radiation R | 105 | 0.8 | 5.515697 | 41.780952 | 40.171429 | 13.780952 |
| Corticospinal tract L | 669 | 5.4 | 7.433486 | 57.043348 | 40.122571 | 24.405082 |
| Corticospinal tract R | 344 | 2.8 | 5.768390 | 34.104651 | 37.162791 | 15.177326 |
| Forceps major | 36 | 0.3 | 3.702282 | 47.722222 | 39.583333 | 44.805556 |
| Inferior fronto-occipital fasciculus L | 37 | 0.3 | 4.688884 | 50.324324 | 21.108108 | 29.027027 |
| Superior longitudinal fasciculus L | 38 | 0.3 | 4.157224 | 58.973684 | 50.421053 | 55.315789 |

### Gray Matter Component (IC2-GM)

The gray matter source map revealed a distributed pattern including regions associated with multiple functional networks (Figure 5.A. Table 2). Positive loadings were observed bilaterally in the inferior and left superior parietal lobule, regions central to the dorsal attention and frontoparietal networks. The temporal lobe showed extensive involvement, including superior, middle, and inferior temporal gyri, areas that form part of the ventral visual stream, semantic processing and social cognition. Frontal contributions included the inferior frontal gyrus and insula, key nodes of the salience networks. Somatosensory regions were represented through increased postcentral gyrus representation. Medial temporal structures including the parahippocampal gyrus, uncus, and fusiform gyrus that are related to memory, visual recognition and emotion. The posterior cingulate cortex (BA31), a hub of the default mode network, also demonstrated positive loadings.

**Figure 5:**
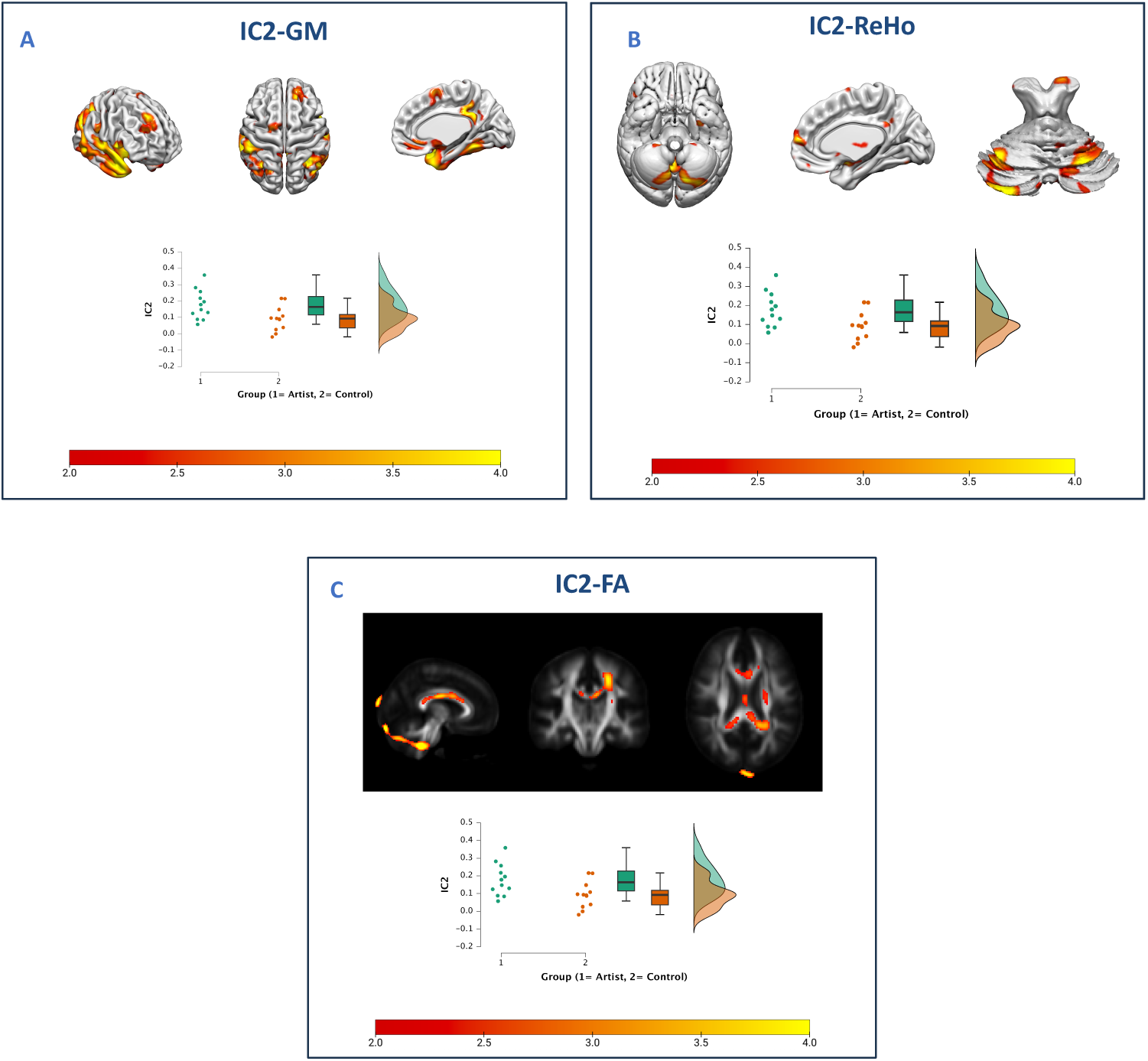
Modality-Specific Spatial Maps and Group Differences for Joint Component IC2. This figure presents the source component loadings for joint Independent Component 2 (IC2), separated by modality: GM (A), ReHo (B), and FA (C). Spatial maps are thresholded at *z* > 2 for visualization purposes only; this threshold is not intended to indicate statistical significance and does not reflect multiple comparison correction. Only positive values were identified across all three modalities. Boxplots below each panel illustrate subject-wise IC2 loading coefficients, showing significantly higher expression in artists compared to controls.

### ReHo Component (IC2-ReHo)

The regional homogeneity map showed increased local functional coherence primarily in subcortical and cerebellar regions (Figure 5.B., Table 2). The basal ganglia structures including lentiform nucleus, putamen, and caudate showed enhanced local synchronization, indicating involvement of motor control and reward processing circuits. Cerebellar regions including the culmen, declive, and pyramis demonstrated increased regional coherence, suggesting engagement of cerebellar networks involved in motor coordination and cognitive processing. In addition, the medial frontal gyrus’s contribution suggests that executive control networks are involved. Moreover, subcortical structures, including the subcallosal gyrus and the claustrum, exhibited positive loadings, showing their association with emotional and integrative processing.

### White Matter Component (IC2-FA)

The fractional anisotropy map identified enhanced structural connectivity in several major white matter pathways (Figure 5.C; Table 3). The anterior thalamic radiations, connecting thalamic nuclei with prefrontal regions, showed increased FA values bilaterally. The corticospinal tracts (bilateral), fundamental for sensorimotor integration and voluntary movements, demonstrated enhanced structural integrity. The forceps major of the corpus callosum (Posterior Forceps), facilitating interhemispheric communication between occipital regions showed increased FA values. Enhanced structural properties were revealed in long-range association fibres, including the left inferior fronto-occipital fasciculus, which connects the prefrontal cortex with the parietal, posterior temporal and occipital cortices, and the left superior longitudinal fasciculus.

### Correlation Analysis Between IC2 Mixing Coefficients and Vividness of Mental Imagery

To further explore the cognitive relevance of the joint component that distinguishes visual artists from non-artists, we examined the relationship between IC2 loading coefficients and Vividness of Visual Imagery Questionnaire (VVIQ; Marks, 1973) scores. Due to incomplete questionnaire data from one participant, the final analysis was conducted on 23 (Total VVIQ: N-1) participants. Pearson correlation revealed a significant negative association between IC2 loadings and total VVIQ scores (*r* = −.467, *p* = .025), indicating that individuals who expressed IC2 more frequently also reported more vivid visual imagery (See Figure 4.). While the correlation was negative, this is consistent with the scoring direction of the VVIQ, whereby lower scores reflect greater vividness. The effect size, computed as Fisher’s z, was –0.506, with a standard error of 0.224. Therefore, this result suggests that greater IC2 expression is associated with enhanced vividness visual imagery ability.

**Tables 2 and 3**. IC2 Modality-Specific Positive Regions and Tracts.

These tables report the positively expressed clusters derived from the source component IC2 for each modality (GM, ReHo, and FA). For Tables 2 and 3, only clusters exceeding a peak Z-score of 4.0 for GM and 3.5 for ReHo and FA, and with volumes greater than 1 cm³, are presented. Coordinates were initially identified in MNI space, then transformed into Talairach space for anatomical labeling. Spatial extent is reported in cubic centimeters (cc), and the peak Z-value along with its corresponding MNI coordinates are indicated for each region.

Regions not matched to the Brodmann Area atlas are marked with an asterisk (*).

## Discussion

In this study, we employed a novel multimodal neuroimaging approach using joint independent component analysis to characterize the neural correlates of professional visual artistic expertise. By integrating gray matter volume, regional homogeneity, and fractional anisotropy data, we identified a multimodal component (IC2) that significantly distinguished professional visual artists from matched controls (*p* = .020, Cohen’s *d* = 1.028). This component revealed coordinated adaptations across structural and functional measures, providing the first comprehensive evidence of how professional artistic expertise manifests across multiple modalities simultaneously.

Our primary finding—that IC2-GM encompasses both traditional creativity networks and domain-specific adaptations—aligns and extends the first hypothesis. The gray matter component revealed structural representations of key DMN and ECN nodes, including the posterior cingulate cortex, insula, and parietal regions, consistent with established models of creative cognition (Beaty et al., 2016). Our findings echo De Pisapia et al. (2016), who observed enhanced DMN-ECN functional connectivity in professional artists during creative planning. While caution is warranted in functionally interpreting ICA-derived networks due to their data-driven nature, prior work demonstrates that spatial patterns from ICA-based decomposition of structural MRI data often recapitulate canonical functional networks including the DMN, ECN, and salience networks (Gupta et al., 2019), supporting their interpretability at the large-scale network level.

The involvement of DMN regions takes on additional significance considering Vessel and colleagues’ findings that the DMN is consistently activated during intense aesthetic experiences, particularly when viewing artworks evoking strong emotional or personal relevance (Vessel et al., 2012, 2019; Belfi et al., 2019). These insights align with our IC2-GM results, potentially reflecting structural underpinnings of deep aesthetic and ideational engagement characteristic of professional artists. The convergence of our structural findings with functional evidence suggests that years of artistic training may produce lasting anatomical adaptations in networks supporting both creative ideation and aesthetic appreciation. Beyond core creativity networks, IC2-GM revealed regions specific to visual artistic expertise. The fusiform gyrus, crucial for face and expression recognition (Adolphs, 2001), showed increased gray matter, consistent with Seabra et al.’s (2022) finding of enhanced N170 amplitude in visual artists. The inferior temporal gyrus, known for high-level visual processing, likely supports semantic visual cognition in artistic ideation (Lin et al., 2020; Davey et al., 2016; Kenett et al., 2023). Notably, our unsupervised approach independently identified Heschl’s gyrus (BA42), previously reported by Grecucci et al. (2023a) as a key GM region distinguishing artists from non-artists, however the directionality effect was uncertain, therefore, current results strengthening evidence for its role in artistic expertise.

The extensive parietal involvement deserves special consideration given its theoretical importance in visual arts. The parietal cortex integrates multisensory information to create spatial frameworks essential for both perceptual analysis and motor execution (Takeuchi et al., 2010a). This function is particularly relevant for the dorsal visual “where” stream, processing object location, movement, and spatial organization crucial for visual artistic thinking (Hong et al., 2023). Our findings build on Jung et al.’s (2010b) work linking the angular gyrus to creative achievement, supported by behavioral evidence that artistically trained individuals excel in spatial ability assessments (Rivero et al., 2024).

The cerebellar findings provide compelling evidence for this structure’s role beyond motor control (Coolidge, 2021). Enhanced regional homogeneity (IC2-ReHo) in the culmen, declive, and pyramis represents, to our knowledge, the first empirical evidence of altered cerebellar functional organization in professional visual artists, addressing the gap identified by Adamaszek et al. (2022). These findings align remarkably with cerebellar-creativity hypothesis, which proposes creativity emerges from blending cerebellar internal models at varying abstraction levels (Coolidge, 2021). The specific cerebellar findings likely reflect growing perceptual proficiency following continuous training in drawing, potentially associated with developments in capturing procedural meaning and enhanced visual perception through the deconstruction and reconstruction of visual scenes that occurs during artistic practice (Adamaszek et al., 2022). According to Ito’s (2008) framework, the cerebellum generates anticipatory and predictive internal models based on sensory information—just as internal models of motor movements generate predictions about sensory outcomes, cerebellar models could generate abstract concepts and thoughts (Coolidge, 2021). In our context, artistic expression may represent the cerebellum transforming thoughts into movements with brushes onto canvas, suggesting the cerebellum treats creative ideation similarly to motor planning. The specific cerebellar regions identified have established connections to both DMN and ECN (Han et al., 2022), potentially explaining the enhanced network interaction observed in artists during creative tasks (De Pisapia et al., 2016). Our functional results align with previous studies that identified the cerebellum’s role in artistic cognition (Makuuchi et al., 2003; Chamberlain et al., 2014).

Our white matter findings revealed enhanced integrity in pathways critical for artistic expertise. Enhanced integrity in the left inferior fronto-occipital fasciculus provides the structural basis for the perceptual and motor advantages that artists have. These advantages largely explain why artists often excel at drawing (Kozbelt, 2001; Kozbelt & Seeley, 2007). Moreover, increased fractional anisotropy (FA) in the bilateral corticospinal tracts reflects the motor expertise necessary for artistic execution.

The correlation between IC2 expression and vividness of visual imagery highlights the neural basis of the “mind’s eye” in artistic creativity. This finding aligns with Morris-Kay’s (2010) evolutionary framework, which positions mental imagery as fundamental to creation of art.

IC2 integrates visuospatial, memory, and executive regions, which support the generation, maintenance, and manipulation of mental images necessary for transforming imagination into artistic expression. This finding is consistent with evidence showing that skilled artists have unique perceptual and cognitive abilities. They excel in visual memory and generating and transforming mental images (Kozbelt & Seeley, 2007; Vodyanyk et al., 2023). Interestingly, both the vividness and the transformativeness of imagery have been positively associated with broader creativity (Jankowska & Karwowski, 2015). Therefore, our results suggest that the neural adaptations reflected in IC2 expression may underpin the enhanced imagery and perceptual capacities that characterize artistic expertise.

It’s important to note that structural adaptations in sensorimotor, visuospatial, and control-related regions have been reported in other high-skill professionals. Musicians show enhanced gray matter in motor and auditory regions (Gaser & Schlaug, 2003), mathematicians in parietal areas for spatial manipulation (Aydin et al., 2007), and pilots and surgeons in frontal-parietal regions for fine motor coordination and decision-making (Ahamed et al., 2014; Karabanov et al., 2019). These parallels suggest our findings may reflect general principles of brain plasticity in response to intensive domain-specific training, therefore results should be interpret considering this fact.

The enhanced regional homogeneity in basal ganglia structures (putamen and caudate) suggests potential dopaminergic system involvement in artistic expertise. Given dopamine’s established role in creativity through motivation, cognitive flexibility, and reward processing (Kenett et al., 2023), and evidence that dopaminergic manipulation influences creative performance (Lanni et al., 2008), these findings open important avenues for future research. This has immediate clinical relevance for understanding creativity in Parkinson’s disease, where dopaminergic dysfunction in the caudate correlates with cognitive changes (Niethammer et al., 2013). Art training-based approaches for PD, already under investigation (Spee et al., 2025), may benefit from understanding these neural mechanisms.

## Limitations

Several limitations warrant consideration. Our sample size of 24 participants, while typical for neuroimaging studies of specialized populations, limits statistical power and generalizability. Sensitivity analysis using G*Power indicated our study could detect large effects (Cohen’s d ≥ 1.20) with 80% power. Our observed effect (d = 1.03) approached this threshold, suggesting a genuine large effect, though this estimate should be interpreted cautiously given the limited sample. The inherent difficulty of recruiting professional artists formally recognized by institutional art systems necessitated this constraint.

Participant recruitment also presents limitations. Visual artists were recruited exclusively through a contemporary art museum, which may have introduced bias toward a certain profile of artists. Meanwhile, the control group was defined only by the absence of visual art experience, without further consideration of their professional or creative background. Future studies could address this by including controls from more specific groups.

To enhance reliability of jICA, we employed ICASSO stability analyses with ten iterations. Our jICA model produced an optimal number of components as calculated by MDL (Li et al., 2007), and consistent with prior ICA-based neuroimaging studies treating component-level inference as low-dimensional (Grecucci et al., 2023b), we did not apply standard multiple comparison corrections. While IC2 (p = 0.020) would not survive stringent Bonferroni correction (0.05/7 = 0.007), given the exploratory nature and data-driven methodology, these findings provide valuable initial insights requiring replication in larger samples.

The cross-sectional design prevents causal inferences about whether observed differences result from training or predisposing factors. Additionally, our focus on visual artistic expertise doesn’t capture creativity across other domains like music, dance, or literature, which likely engage partially overlapping but distinct neural systems. We also lacked behavioral or psychometric assessments of artistic creativity beyond the VVIQ. While visual imagery correlates with creative thinking dimensions (Palmiero et al., 2011), it represents only one aspect of artistic expertise.

## Conclusions

Using joint independent component analysis, we identified a multimodal component that differentiated artists from controls and correlated with visual imagery vividness. This component (IC2) revealed coordinated adaptations across cerebellar-cortical circuits, basal ganglia structures, major white matter pathways, and key DMN-ECN nodes, demonstrating that artistic expertise emerges from the coordinated adaptation of multiple neural systems. Our findings advance understanding of artistic creativity by showing that professional expertise extends beyond traditional creativity networks to encompass cerebellar, sensorimotor, and subcortical systems. These results have implications for education, clinical practice, and understanding human creativity as a core evolutionary adaptation. Future research should investigate other artistic domains, and integrate comprehensive behavioral assessments to fully understand the relationship between neural architecture and creative performance.

## Declarations

### Funding

This study was funded by a grant titled “Capolavori della mente: creatività, cognizione ed emozione nelle opere delle collezioni MART” from the Fondazione Caritro (Cassa di Risparmio di Trento e Rovereto) awarded to F.B/D.M.

**Conflicts of interest/Competing interests** Authors declare no conflict of interest **Ethics approval**

This study adhered to the ethical standards of the Declaration of Helsinki and was approved by the Ethics Committee of the University of Trento. Protocol number: 2008006 granted to Prof. D.M. project entitled “Visual perception, art and the brain”. Web page of the Committee: https://www.unitn.it/en/research/responsible-research/ethics-and-integrity/research-ethics-committee

### Consent to participate

Participants provide a signed informed consent to participate to the study

### Availability of data and materials

Data will be available upon reasonable request

### Author Contributions

*ET and AG:* methodology, data curation, formal analysis, and writing—original draft preparation, Writing – review & editing.*NDP, DM, FB:* conceptualization, data acquisition, Writing – review & editing.

