## Supplementary_Material for "The Artists’ Brain: A Data Fusion Approach to Characterize the Neural Bases of Professional Visual Artists"

| **Table 4. Participant Demographics and Imagery Vividness Scores** | | | |
| --- | --- | --- | --- |
| **Group** | **Age** | **Sex** | **VVIQ (total score)** |
| ARTIST | 31,63 | M | 36 |
| ARTIST | 22,44 | M | 19 |
| ARTIST | 30,77 | M | 28 |
| ARTIST | 23,34 | M | 30 |
| ARTIST | 36,44 | F | 21 |
| ARTIST | 32,65 | M | 36 |
| ARTIST | 37,08 | F | 34 |
| ARTIST | 29,32 | M | 18 |
| ARTIST | 30,18 | M | 38 |
| ARTIST | 36,43 | F | 29 |
| ARTIST | 27,68 | F | 32 |
| ARTIST | 32,85 | M | MISSING |
| CTRL | 23,96 | F | 35 |
| CTRL | 41,23 | M | 44 |
| CTRL | 36,32 | F | 20 |
| CTRL | 29,84 | M | 41 |
| CTRL | 32,49 | M | 35 |
| CTRL | 24,01 | F | 35 |
| CTRL | 28,96 | F | 43 |
| CTRL | 25,42 | M | 33 |
| CTRL | 29,69 | F | 31 |
| CTRL | 32,82 | M | 36 |
| CTRL | 24,14 | F | 44 |
| CTRL | 28,3 | M | 41 |

**Supplementary Material**

**The Artists’ Brain: A Data Fusion Approach to Characterize the Neural Bases of Professional Visual Artists**

**Github Links and Explanations**

- For the FA modality, visualization and tract-level quantification of significant joint components were carried out using a custom FSL-based pipeline developed by Authors and implemented in a UNIX environment (<https://github.com/erdemtaskiran/JHU-tract-component-overlap-FA>)
- All diffusion-weighted imaging data underwent comprehensive preprocessing using the FMRIB Software Library (FSL version 6.0; Jenkinson et al., 2012). The preprocessing pipeline was an automated bash script created by the Authors (<https://github.com/erdemtaskiran/DTI/blob/main/DTI_preprocessing_UNIX>) to ensure consistency across all subjects.

****Visualization of Joint Components****

The joint source matrix (S), obtained from the ICA decomposition, was used to visualize the spatial maps of the significant components for each modality: grey matter (GM), regional homogeneity (ReHo) and fractional anisotropy (FA). Each row of the source matrix, corresponding to a component, was converted to a Z-score map by dividing voxel-wise values by their standard deviation. These normalized Z-maps were then reshaped into 3D brain volumes for each modality.

For GM and ReHo, a threshold of Z > 4.0 and Z > 3.5 respectively was applied to identify regions. These thresholded Z-maps were then converted from MNI to Talairach coordinates using the Group ICA of fMRI Toolbox (GIFT), and anatomical labels were assigned using the Talairach Daemon. The identified brain regions were then visualized using Surf Ice (https://www.nitrc.org/projects/surfice/), and the volume of each region (L/R) was reported in cubic centimeters (cc). Only positive activations were included in the final visualization because no regions showing decreased values exceeded the threshold of Z >- 1.5.

For the FA modality, visualization and tract-level quantification of significant joint components were carried out using a custom FSL-based pipeline developed by Authors and implemented in a UNIX environment (<https://github.com/erdemtaskiran/JHU-tract-component-overlap-FA>). Z-maps corresponding to the significant component were first converted from Analyze (.img/.hdr) to NIfTI (.nii) format using Mango ([https://mangoviewer.com](https://mangoviewer.com/)). These NIfTI files were then thresholded at Z >3.5 and binarized using fslmaths. Only the positive tail of the distribution was retained, as no negatively contributing regions exceeded the threshold (i.e., Z < –3.5), and thus did not meet criteria for inclusion in the final analysis.

To assess the anatomical distribution of the FA component, the resulting binary mask was intersected with each of the 20 major white matter tract masks defined in the JHU ICBM-DTI-81 20-Tract Atlas (2 mm resolution). For each tract, a binary mask was generated and combined with the FA component mask using fslmaths to compute voxel-wise intersections. The number of overlapping voxels was extracted using fslstats. Given that each voxel has a volume of 8 mm³ (2 mm x 2 mm x 2 mm), the total overlap volume was computed and converted into cubic centimeters (1 cm³ = 1000 mm³) for reporting.

To further quantify the component’s spatial distribution, we calculated the percentage of each tract involved using the formula:

*Overlap = (Number of overlapping voxels / Total tract voxels) x 100*

In addition, we extracted the maximum Z-score within each overlapping region to identify peak component intensity and computed the center of gravity (COG) coordinates using the FSL separately. COG provides a single, representative coordinate that captures the spatial center of the component's expression by representing the weighted average location of all activated voxels within a tract. This makes it easy to position the component in the correct place in the body, even when activation is spread across the tract.

Normalization of All Modalities for Multimodal Fusion Analysis

Following the procedure described in Calhoun et al. (2006a), all modalities (GM, FA, ReHo) were resampled to a voxel size of (2 mm) ³ to ensure consistent spatial resolution across modalities prior to fusion analysis. The resampling was performed using a custom automated Bash pipeline leveraging FSL’s FLIRT tool with trilinear interpolation, using the sform/qform matrices to preserve spatial alignment. Post-processing verification ensured all outputs were correctly resliced and free of corruption. After feature extraction, the 3D brain image of each participant was reshaped into a one-dimensional non-zero voxel vector and stacked to form a matrix with dimensions of 24 x [number of voxels] for each imaging modality. The feature matrix for each modality was then normalized to have the same average sum-of-squares across all participants and voxels within that modality, ensuring the raw data, which exhibited different ranges across modalities, were scaled comparably for joint analysis. A single normalization factor was used for each data type; thus, following normalization, the relative scaling within each modality was preserved while units between data types were matched in a least-squares sense. Normalization was performed on group level, so covariations among subjects were preserved (Wang et al., 2015).
